# Gene expression effects of lithium and valproic acid in a serotonergic cell line

**DOI:** 10.1101/227652

**Authors:** Diana Balasubramanian, John F. Pearson, Martin A. Kennedy

**Affiliations:** Gene Structure and Function Laboratory and Carney Centre for Pharmacogenomics, Department of Pathology, University of Otago, Christchurch, New Zealand; Biostatistics and Computational Biology unit, University of Otago, Christchurch, New Zealand

**Keywords:** RNA-Seq, valproic acid, gene expression, mood stabilizer, pharmacogenomics

## Abstract

Valproic acid (VPA) and lithium are widely used in the treatment of bipolar disorder. However, the underlying mechanism of action of these drugs is not clearly understood. We used RNA-Seq analysis to examine the global profile of gene expression in a rat serotonergic cell line (RN46A) after exposure to these two mood stabilizer drugs. Numerous genes were differentially regulated in response to VPA (log_2_ fold change ≥ 1.0; i.e. odds ratio of ≥ 2, at FDR <5%), but only two genes (*Dynlrb2* and *Cdyl2*) showed significant differential regulation after exposure of the cells to lithium, with the same analysis criteria. Both of these genes were also regulated by VPA. Many of the differentially expressed genes had functions of potential relevance to mood disorders or their treatment, such as several serpin family genes (including neuroserpin), *Nts* (neurotensin), *Maob* (monoamine oxidase B) and *Ap2b1*, which is important for synaptic vesicle function. Pathway analysis revealed significant enrichment of Gene Ontology terms such as extracellular matrix (ECM) remodelling, cell adhesion and chemotaxis. This study in a cell line derived from the raphe nucleus has identified a range of genes and pathways that provide novel insights into the therapeutic action of the commonly used mood stabilizer drugs.

## Introduction

Valproic acid (VPA) and lithium are two of the most widely used mood stabilizers and are regularly prescribed as first line therapy in bipolar disorder [1]. They are also used as combination therapy in patients who fail to respond to monotherapy with either drug [2]. The mechanisms involved in the therapeutic action of these drugs in mood stabilization remain poorly understood. The two drugs have highly dissimilar chemical structures yet seem to have similar efficacy in bipolar patients [3] Lithium is known to exert a strong effect on a number of neuronal signalling pathways. Of these, inhibition of inositol monophosphatase (IMPase) thereby inhibiting the inositol pathway and the glycogen synthase kinase-3 β (GSK-3β) signalling pathway, is considered a primary direct target of lithium [4-7]. VPA, on the other hand, is a histone deacetylase (HDAC) inhibitor which regulates a number of genes involved in various signalling pathways, including indirect modulation of GSK-3 and the inositol pathways [8-10].

Although no single molecule has been identified as a common direct target of both VPA and lithium, a number of signalling pathways and molecules have been studied as shared indirect targets between the two structurally different mood drugs [11-17].

In this study, we used RNA-Seq analysis to examine the global profile of gene expression in the rat serotonergic RN46A cell line, in response to VPA and lithium. Dysregulation of the serotonergic system has long been implicated in mood disorders [18-20]. Serotonin in the brain is released from serotonergic neurons originating from the dorsal and median raphe nuclei in the mid-brain [21, 22]. The RN46A cell line used in this study was derived by retroviral transduction of a cell from embryonic rat medullary raphe nucleus [23]. This neuronally restricted cell line was chosen due to its serotonergic nature, and even in the relatively undifferentiated state used in this study, the cells express both the serotonin transporter (SERT) and the high-affinity serotonin receptor, 5HT-1A as well as low levels of tryptophan hydroxylase [24, 25]. Previously, we have used this cell line for candidate gene expression studies in response to various antidepressants [26-28].

## Materials and Methods

### Cell culture and RNA extraction

RN46A cells [23] were cultured in Dulbecco’s Modified Eagle Medium: (DMEM/F12 with GlutaMAX™-I) and supplemented with 5% FBS and 250 μg/ml Geneticin^®^ (G418). Cells were exposed to 0.5 mM VPA or lithium chloride (LiCl) for 72 h. VPA as sodium valproate and lithium chloride (LiCl) were purchased from Sigma Aldrich (St. Louis, MO, USA). Stock solutions of the drugs were prepared in ultrapure Milli-Q^®^ (MQ) water (Millipore, MA, USA) and diluted in the culture medium to obtain the required concentration of 0.5 mM. Untreated cells cultured for 72 h were used as control. Total RNA was isolated from the cells using Trizol^®^ LS Reagent (Invitrogen, Carlsbad, CA). The quality of the total RNA extracted was analysed using the MultiNA MCE^®^-202-microchip electrophoresis system (Shimadzu Corporation, Kyoto, Japan). The DNase-I treated RNA samples were then stored in RNAstable (Biomatrica, San Diego, CA, USA) and shipped to Otogenetics Corporation (Norcross, GA, USA) at ambient temperature for RNA-Seq. The drug exposure experiment was repeated twice, and RNA samples from both experiments were prepared and sent for RNA-Seq analysis.

### RNA-Seq analysis

RNA-Seq was performed on the Illumina HiSeq™ 2000 sequencing system (Illumina Inc. San Diego, CA, USA) by Otogenetics Corporation (Norcross, GA, USA). 100 bp paired-end sequencing generated >20 million reads per sample, which were mapped to the Rat rn4 reference genome (UCSC Baylor3.4/rn4; Nov 2004). For read mapping, a cloud based sequencing data analysis platform provided by DNAnexus (DNAnexus Inc., Mountain View, CA, USA) was used. DNAnexus uses a Bayesian technique for a “probabilistic approach” to map reads to the genome. Reads are mapped to a location when the posterior probability of mapping at that location is ≥0.9 (≥ 90%). These posterior probabilities are then summed to generate the read counts for each gene [29, 30].

For analysis of differential gene expression between untreated cells and cells treated with lithium or VPA, we used the R (Vienna, Austria) package DESeq2 v 1.18.0 [31, 32].

DESeq2, like its predecessor DESeq [33], uses a negative binomial distribution model to test for differential expression from the read count data, while maintaining control over the type-I error rate. Shrinkage estimators are used for fold changes and dispersions for increased stability and reproducibility of analysis [31, 32]. Differential expression for each gene was reported as a fold change along with the statistical significance (p-values). Also reported are the p-values adjusted for multiple testing using the Benjamini-Hochberg procedure to control for false discovery rate (FDR) [34].

### Gene ontology studies

Metacore™ pathway analysis software (Thompson Reuters, USA, USA) was used for analysis of differentially expressed genes. Metacore™ is an integrated software tool based on a manually curated database of metabolic and signalling pathways, transcription factors, and protein-DNA/protein-RNA/protein-protein interactions. It allows functional analysis of microarray or RNA-Seq data and includes Gene Ontology (GO)/processes enrichment analysis, biomarker assessment and drug target/toxicity networks [35]

The Metacore™ database provides several different workflows for the analysis of a given dataset. These include enrichment analysis, toxicity analysis and biomarker assessment workflows. In this study, the enrichment analysis workflow was used. This shows the most enriched ontologies or pathway maps in the gene lists uploaded to Metacore™. This analysis mainly includes GO ontologies and KEGG pathways apart from the proprietary Metacore pathways which are generated at Metacore™ from the various (~1000) canonical pathways. The biomarker assessment workflow which helps to analyse datasets focussing on specific disorders was used for analysis of our RNA-Seq dataset. The disease category ‘mental disorders’ was used to generate category-specific pathway maps.

### Quantitative real time PCR

Quantitative real time PCR (qPCR) was performed on the LightCycler^®^ 480 System (Roche Applied Science, Mannheim, Germany). The Universal Probe Library (UPL) System (Roche Applied Science) was used for relative quantification assays and primers for each candidate gene were designed using the UPL assay design centre [36]. cDNA synthesis was carried out using Superscript™ III first strand synthesis system (Invitrogen, Carlsbad, CA) according to the manufacturer’s instructions. All reactions were run in triplicate and gene expression differences were measured using normalization to three reference genes (*Actb*, *G6pd*, and *Rnf4*) as previously used in this laboratory [37] A calibrator cDNA sample obtained from untreated RN46A cells was used in each run to normalize for inter-run variation. The primer sequences and UPL probes used in this assay are listed in Supplementary Table 1.

For selected genes, absolute quantification was performed using the chip-based Quantstudio 3D digital PCR system (Thermofisher Scientific). For this, ~10ng/uL of cDNA, synthesized as above, was mixed with 2X QuantStudio™ 3D Digital PCR 20K Chip mastermix, 300 nM of primers (forward and reverse), and appropriate volume of MQ water and loaded onto the chips using the Quantstudio chip loader. PCR was performed on the ProFlex™ 2x Flat PCR System (ThermoFischer Scientific) using the following cycling parameters: 96□°C for 10□minutes, 39 cycles of 60□°C for 2□min and 98□°C for 30□sec, and a final extension at 60□°C for 2□min. The chips were read using the QuantStudio 3D Digital PCR instrument and analysed using the QuantStudio 3D AnalysisSuite™ Cloud Software.

## Results

### Analysis of differential gene expression in RN46A cells in response to VPA and lithium

In this study, we used RNA-Seq to investigate the effects of VPA and lithium treatment on global gene expression in RN46A cells. RN46A cells were exposed to either 0.5 mM VPA or 0.5 mM lithium for 72 h, as previously described [37]. A total of 16,453 genes were analysed for differential expression using the R-package DESeq2 v1.10.1

Genes showing a log_2_ fold change of ≥1.0-fold (i.e. odds ratio ≥ 2) at an FDR of <5% were considered to be significantly regulated. Using these criteria, a total of 88 genes were found to be differentially regulated by VPA, with 70 genes upregulated (≥ log_2_1.0-fold) and 18 genes downregulated (≤ log_2_1.0-fold). In comparison, lithium exposure resulted in differential expression of only 2 genes, *Dynlrb2* (upregulated 3.2-fold, FDR 1.3×10^21^) and *Cdyl2* (downregulated −2.3fold, FDR 2.26×10^9^. Both these genes were also regulated by VPA in the same direction (Tables 1 and 2).

**Table 1.**
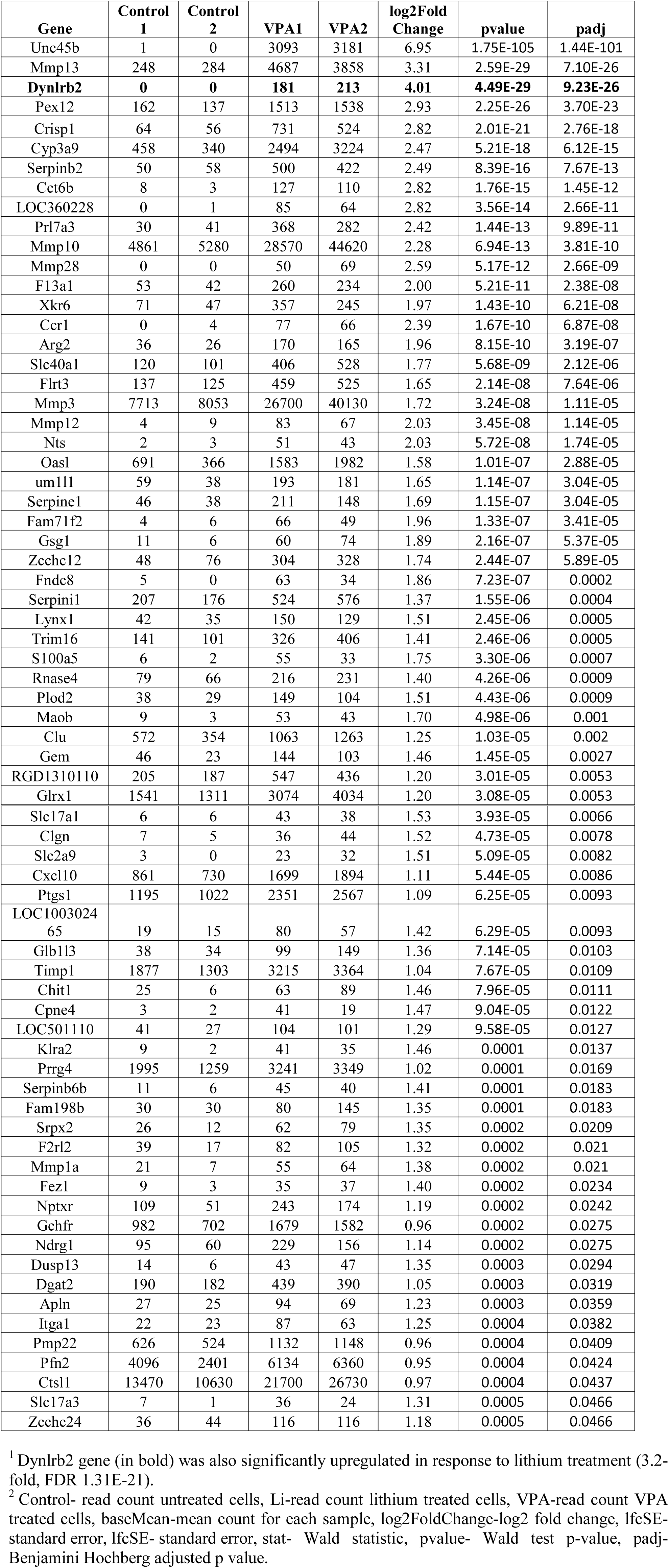
List of genes significantly upregulated in response to VPA (Log_2_ fold change of ≥1.0-fold, FDR<0.05).

**Table 2.** List of genes significantly downregulated in response to VPA (Log2 fold change of≥ 1.0-fold, FDR<0.05).

| Gene | Control 1 | Control 2 | VPA1 | VPA2 | log2Fold Change | pvalue | padj |
| --- | --- | --- | --- | --- | --- | --- | --- |
| Ap2b1 | 1780 | 1735 | 0 | 1 | -6.02 | 3.55E-74 | 1.46E-70 |
| Slfn3 | 1062 | 1495 | 104 | 84 | -2.75 | 2.06E-17 | 2.11E-14 |
| Heatr6 | 163 | 74 | 4 | 3 | -2.66 | 2.89E-13 | 1.83E-10 |
| Taf15 | 79 | 79 | 1 | 0 | -2.71 | 4.12E-13 | 2.42E-10 |
| Slfn8 | 3621 | 3798 | 639 | 694 | -1.93 | 3.08E-11 | 1.49E-08 |
| Nle1 | 251 | 29 | 0 | 1 | -2.06 | 3.8E-08 | 1.2E-05 |
| **Cdyl2** | **86** | **67** | **7** | **8** | **-1.94** | **8.31E-08** | **2.44E-05** |
| Eno3 | 1483 | 935 | 351 | 420 | -1.22 | 1.78E-05 | 0.003 |
| Lig3 | 314 | 95 | 10 | 34 | -1.59 | 2E-05 | 0.004 |
| Rad51l3 | 67 | 33 | 0 | 6 | -1.51 | 5.75E-05 | 0.009 |
| Kdelc2 | 309 | 431 | 91 | 106 | -1.33 | 6.35E-05 | 0.009 |
| Dhrs4 | 201 | 85 | 35 | 30 | -1.35 | 8.39E-05 | 0.011 |
| Cxcl3 | 48 | 55 | 3 | 10 | -1.41 | 0.0002 | 0.021 |
| Csf3 | 100 | 63 | 14 | 22 | -1.30 | 0.0003 | 0.028 |
| Igtp | 310 | 244 | 74 | 108 | -1.11 | 0.0004 | 0.044 |
| Penk | 968 | 618 | 342 | 307 | -0.95 | 0.0005 | 0.047 |
| Ifi44 | 34 | 26 | 4 | 3 | -1.31 | 0.0005 | 0.048 |
| Ccnd1 | 4463 | 2665 | 1686 | 1072 | -0.99 | 0.0005 | 0.051 |
| Hspb6 | 1520 | 840 | 354 | 483 | -1.03 | 0.0006 | 0.058 |
| Mxra8 | 1084 | 331 | 216 | 176 | -1.15 | 0.0007 | 0.065 |

<sup>1</sup>Cdyl2 gene (highlighted) was also significantly downregulated in response to lithium treatment (-2.3fold, FDR 2.26E-09).
<sup>2</sup>Control- read count untreated cells, Li-read count lithium treated cells, VPA-read count VPA treated cells, baseMean-mean count for each sample, log2FoldChange-log2 fold change, lfcSE-standard error, lfcSE- standard error, stat- Wald statistic, pvalue- Wald test p-value, padj-Benjamini Hochberg adjusted p value.

### Functional annotation of RNA-Seq data

The lists of differentially expressed genes from the VPA RNA-Seq dataset (log_2_ fold change ≥1.0-fold, FDR< 5%) were examined using Metacore™ pathway analysis software (Thompson Reuters, USA, formerly GeneGo inc., USA). Enrichment analysis workflow was used, which generated Metacore pathway maps and highlighted several GO processes. The GO processes and Metacore pathways enriched after VPA treatment are summarized in Tables 3 and 4.

**Table 3.**
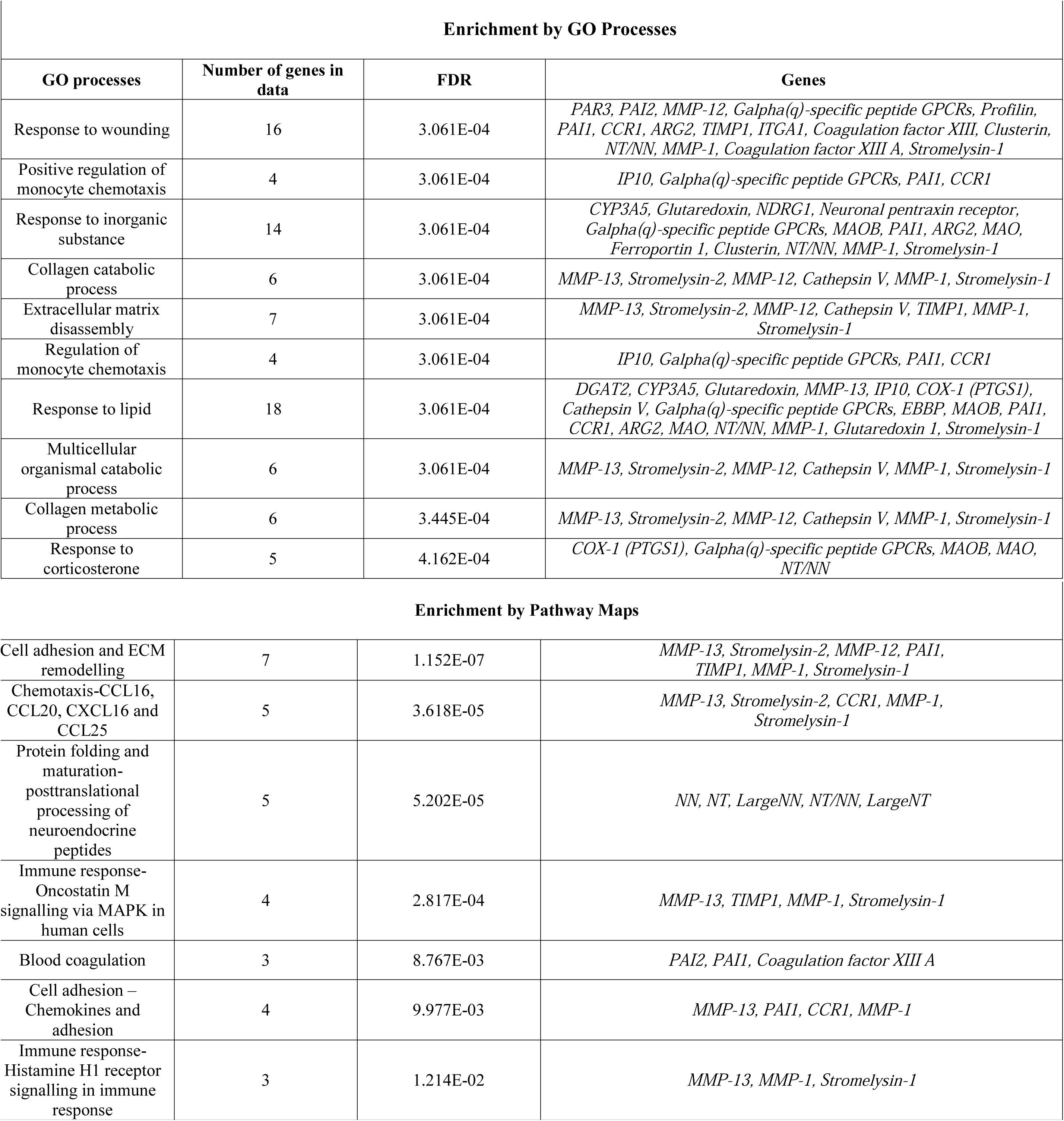
Enrichment analysis for genes upregulated in response to VPA

| **Enrichment by GO Processes** |  |  |  |
| --- | --- | --- | --- |
| **GO processes** | **Number of genes in data** | **FDR** | **Genes** |
| Response to wounding | 16 | 3.061E-04 | *PAR3, PAI2, MMP-12, Galpha(q)-specific peptide GPCRs, Profilin, PAII, CCR1, ARG2, TIMP1, ITGA1, Coagulation factor XIII, Clusterin, NT/NN, MMP-1, Coagulation factor XIII A, Stromelysin-1* |
| Positive regulation of monocyte chemotaxis | 4 | 3.061E-04 | *IP10, Galpha(q)-specific peptide GPCRs, PAII, CCR1* |
| Response to inorganic substance | 14 | 3.061E-04 | *CYP3A5, Glutaredoxin, NDRG1, Neuronal pentraxin receptor, Galpha(q)-specific peptide GPCRs, MAOB, PAII, ARG2, MAO, Ferroportin 1, Clusterin, NT/NN, MMP-1, Stromelysin-1* |
| Collagen catabolic process | 6 | 3.061E-04 | *MMP-13, Stromelysin-2, MMP-12, Cathepsin V, MMP-1, Stromelysin-1* |
| Extracellular matrix disassembly | 7 | 3.061E-04 | *MMP-13, Stromelysin-2, MMP-12, Cathepsin V, TIMP1, MMP-1, Stromelysin-1* |
| Regulation of monocyte chemotaxis | 4 | 3.061E-04 | *IP10, Galpha(q)-specific peptide GPCRs, PAII, CCR1* |
| Response to lipid | 18 | 3.061E-04 | *DGAT2, CYP3A5, Glutaredoxin, MMP-13, IP10, COX-1 (PTGS1), Cathepsin V, Galpha(q)-specific peptide GPCRs, EBBP, MAOB, PAII, CCR1, ARG2, MAO, NT/NN, MMP-1, Glutaredoxin 1, Stromelysin-1* |
| Multicellular organismal catabolic process | 6 | 3.061E-04 | *MMP-13, Stromelysin-2, MMP-12, Cathepsin V, MMP-1, Stromelysin-1* |
| Collagen metabolic process | 6 | 3.445E-04 | *MMP-13, Stromelysin-2, MMP-12, Cathepsin V, MMP-1, Stromelysin-1* |
| Response to corticosterone | 5 | 4.162E-04 | *COX-1 (PTGS1), Galpha(q)-specific peptide GPCRs, MAOB, MAO, NT/NN* |

| Enrichment by Pathway Maps |  |  |  |
| --- | --- | --- | --- |
| Cell adhesion and ECM remodelling | 7 | 1.152E-07 | *MMP-13, Stromelysin-2, MMP-12, PAI1, TIMP1, MMP-1, Stromelysin-1* |
| Chemotaxis-CCL16, CCL20, CXCL16 and CCL25 | 5 | 3.618E-05 | *MMP-13, Stromelysin-2, CCR1, MMP-1, Stromelysin-1* |
| Protein folding and maturation-posttranslational processing of neuroendocrine peptides | 5 | 5.202E-05 | *NN, NT, LargeNN, NT/NN, LargeNT* |
| Immune response-Oncostatin M signalling via MAPK in human cells | 4 | 2.817E-04 | *MMP-13, TIMP1, MMP-1, Stromelysin-1* |
| Blood coagulation | 3 | 8.767E-03 | *PAI2, PAI1, Coagulation factor XIII A* |
| Cell adhesion – Chemokines and adhesion | 4 | 9.977E-03 | *MMP-13, PAI1, CCR1, MMP-1* |
| Immune response-Histamine H1 receptor signalling in immune response | 3 | 1.214E-02 | *MMP-13, MMP-1, Stromelysin-1* |

**Table 4.** Enrichment analysis for genes downregulated in response to VPA

| Enrichment by GO Processes |  |  |  |
| --- | --- | --- | --- |
| GO processes | Number of genes in data | FDR | Genes |
| Multi-organism processes | 10 | 1.027E-01 | *ENO, Beta-adaptin 2, GRO-1, G-CSF, GRO-3, ENO3, IGTP, TAFs, IFI44, Enkephalin A* |
| Response to external biotic stimulus | 6 | 1.027E-01 | *ENO, GRO-1, G-CSF, GRO-3, IGTP, IFI44* |
| Skeletal muscle tissue regeneration | 2 | 1.027E-01 | *ENO, ENO3* |
| Gluconeogenesis | 2 | 1.027E-01 | *ENO, ENO3* |
| Chemokine-mediated signalling pathway | 2 | 1.027E-01 | *GRO-1, GRO-3* |
| Hexose biosynthetic process | 2 | 1.027E-01 | *ENO, ENO3* |
| Cellular response to lipopolysaccharide | 3 | 1.027E-01 | *GRO-1, G-CSF, GRO-3* |
| Enrichment by Pathway Maps |  |  |  |
| Glycolysis and gluconeogenesis p.3/Human version | 7 | 1.152E-07 | *MMP-13, Stromelysin-2, MMP-12, PAI1, TIMP1, MMP-1, Stromelysin-1* |
| Th17 cells in CF | 5 | 2.808E-04 | *MMP-13, TIMP1, MMP-1, Stromelysin-1* |
| Immune response-IL7 signalling pathways | 4 | 2.808E-04 | *MMP-13, TIMP1, MMP-1, Stromelysin-1* |
| Development-role of nicotinamide in GCSF-induced granulopoiesis | 4 | 9.977E-03 | *MMP-13, PAI1, CCR1, MMP-1* |
| Development –Role of G-CSF in | 3 | 1.214E-02 | *MMP-13, MMP-1, Stromelysin-1* |
| hematopoietic stem cell<br>mobilization |  |  |  |

### Validation of RNA-Seq results using qPCR

Eight genes identified by RNA-Seq to be differentially expressed were chosen for qPCR validation (Table 5). Genes for validation were selected on the basis of significant differential expression in response to either or both the drugs.

**Table 5.** Validation of RNA-Seq results using qPCR

| Gene | RNA-Seq |  |  |  | qPCR validation |  |  |  |
| --- | --- | --- | --- | --- | --- | --- | --- | --- |
|  | VPA |  | Lithium |  | VPA |  | Lithium |  |
|  | log<sub>2</sub> Fold change | Adjusted p-value | Fold change | Adjusted p-value | log<sub>2</sub> Fold change | p-value | Fold change | p-value |
| *Ap2b1* | -6.02 | 1.46E-70 | - | - | -0.34 | 0.004 | - | - |
| *Cdyl2* | -1.93 | 2.44E-05 | -2.31 | 2.26E-09 | -0.71 | 0.09 | -4.3 | 0.08 |
| *Dnlrb2\** | 4.01 | 9.23E-26 | 3.27 | 1.31E-21 | Absolute quantification<br>Untreated: 8.8 copies/uL<br>VPA: 210 copies/uL<br>Lithium: 92 copies/uL |  |  |  |
| *Dusp13* | 1.35 | 0.029 | - | - | 1.03 | 0.004 | - | - |
| *Mmp13* | 3.31 | 7.10E-26 | - | - | 1.92 | 0.04 | - | - |
| *Nle1\** | -2.05 | 1.20E-05 | - | - | Absolute quantification<br>Untreated: 506 copies/uL<br>VPA: 3.5 copies/uL |  |  |  |
| *Unc45b* | 6.95 | 1.44E-101 | - | - | 3.48 | 0.06 | - | - |
| *Zcchc12* | 1.74 | 5.89E-05 | - | - | 1.07 | 0.03 | - | - |

\*Absolute quantification for *Dnlrb2* and *Nle1* was carried out using the Quantstudio 3D digital PCR platform (Thermofisher Scientific).

All eight genes showed similar regulation between the two platforms. However, the magnitude of the change differed for some of the genes (Table 5). This was especially true for one of the genes, *Ap2b1*, which was strongly downregulated in response to VPA (log_2_ fold change −6-fold, *p*=1.46×10^-70^) in the RNA-Seq analysis. qPCR data for *Ap2b1* showed a significant but much lesser downregulation (log_2_ fold change of −0.3-fold; *p*<0.005).

For two of the genes, *Dynlrb2* and *Nle1*, absolute quantification was carried out using digital PCR (QuantStudio 3D, ThermoFisher) because these genes were expressed at a level below the detection limits of qPCR assay for relative quantification.

## Discussion

In this RNA-Seq study, we explored the regulation of gene expression by two widely used mood stabilizer drugs, VPA and lithium, in a rat serotonergic cell line (RN46A) originally derived from the medullary raphe nucleus, a central nervous system region strongly implicated in mood disorders and their treatment [38, 39]. This cell line, to a degree, recapitulates the biology of the raphe nucleus, and provide a relevant and tractable *in vitro* model for analysis of drug-induced gene expression changes.

The study revealed expression differences in multiple genes in response to VPA or lithium exposure in RN46A cells, with VPA showing more extensive effects than lithium. A total of 88 genes were significantly differentially expressed after VPA treatment. In comparison, lithium treatment resulted in significant differential expression of only two genes (Tables 1 and 2). The broader differential gene expression effects of VPA were expected, as it is a strong HDAC inhibitor and a potent drug with a wider range of clinical benefits [40-43] as well as some significant side effects, including teratogenicity [44, 45].

We used qPCR and chip-based digital PCR to assess the validity of the differential expression observed in our RNA-Seq dataset. This validation was carried out for a set of the most differentially expressed genes in response to either or both the drugs. Digital PCR was used for two genes which showed low levels of expression. We observed a generally positive correlation with the majority of genes being regulated in the same direction in both platforms with a comparable fold difference.

Bioinformatic tools such as GO analysis or canonical pathway mapping can help to interpret the biological significance of gene expression profiling studies. We used Metacore™ pathway analysis software for functional annotation of the RNA-Seq dataset. Analysis of genes upregulated in response to VPA highlighted biological pathways involved in processes such as the extracellular matrix (ECM) remodelling, cell adhesion and chemotaxis. VPA (0.3-1.2 mM) has previously been shown to regulate cell adhesion molecules such as neuroligin-1 as well as some extracellular matrices in primary rat astrocytes [46]. Interestingly, several matrix metalloproteinase (MMP) family proteases (*Mmp13*, *Mmp10*, *Mmp12*, and *Mmp28*) were significantly upregulated by VPA. MMPs play an important role in both neurogenesis and neuroinflammation, processes that are associated with various neuropsychiatric conditions [47, 48]. Epigenetic mechanisms involved in ECM remodelling have recently been described in melanoma cells [49], Significantly enriched GO terms highlighted include ‘wound healing’, a complex process in which VPA has been previously implicated [50] as well as collagen metabolic/catabolic processes. The MMP family of proteins are known to be involved in these processes [51, 52]. Analysis of genes downregulated by VPA highlighted GO terms such as glycolysis and gluconeogenesis as well as immune response signalling pathways. Taken together, the GO and pathway analyses in this study were consistent with what is known about the action of these drugs, and the biology of mood disorder treatment.

There do not appear to be any studies using RNA-Seq to examine VPA or lithium effects on cell culture or animal tissues, especially in a neuronal setting. However, several microarray studies using VPA and lithium have been performed in the past in search of potential mechanisms of action of these drugs. All these studies have generated lists of differentially expressed genes which could not always be replicated by other studies using similar concentrations of VPA or lithium [53-57]. However, there are some overlaps between our analysis and previous microarray studies using these two drugs in various experimental systems. For example, microarray analysis in ES cells treated with 1 mM VPA for 8 h [58] and 0.5 mM for 4 h [59] showed that a small percentage of genes in these cells are regulated by VPA. Of these the *Ccnd1* gene was common between the above two studies and ours, being downregulated in all cases. Interestingly, a previous study from our laboratory also observed downregulation of *Ccnd1* in rat hippocampus in response to chronic doses of the antidepressant paroxetine [27]. Similar overlaps were also found with microarray studies using lithium in cell culture as well as animal models [56, 60].

Two genes in our study that were significantly regulated by lithium were also coregulated by VPA. *Dynrlb2* was found to be upregulated by both drugs (VPA-log_2_ fold change of 4.01-fold; lithium-log_2_ fold change of 3.2-fold). This gene encodes the Dynein, Light Chain, Roadblock-Type 2 protein. Dynein proteins are one of the major families of cytoskeleton motor proteins in eukaryotes. These proteins use ATP to drive intracellular transport through the microtubule network. A variety of proteins, organelles and mRNAs involved in numerous cellular and developmental processes are transported by dynein proteins [61]. *Cdyl2*, encoding a chromodomain protein, was also markedly downregulated by both the drugs. Chromodomains are considered to be ‘readers’ of the histone methyl-lysine code and are known to regulate transcription in response to epigenetic cues [62, 63]. Neither *Dnrlb2* nor *Cdyl2* have been implicated in the aetiology or treatment of neuropsychiatric disorders, but their functional characteristics and co-regulation by both drugs suggest that further examination of these may be warranted.

This analysis also highlighted several other genes with potentially interesting and relevant functions that were regulated by exposure to VPA. For example, VPA significantly upregulated *Nts* (neurotensin), which has been previously shown to be markedly upregulated by lithium [64], although we did not detect a significant effect on this gene due to lithium in our model system. *Maob*, encoding a subunit of monoamine oxidase, also showed upregulation after VPA exposure. Monoamine oxidase is involved in the metabolism of neurotransmitters such as dopamine and serotonin, and it is a target for several drugs including some antidepressants [65]. Several serine protease inhibitor (serpin) family members were upregulated by VPA, including *Serpinb2*, *Serpine1*, *Serpini1* and *Serpinb6b* (Table 1). Of these serpins, which are involved in many regulatory processes, SerpinI1 (neuroserpin) is primarily secreted by axons in the brain, and may play a role in axonal growth and synaptic plasticity [66]. In humans, mutation in *SERPINI1* lead to childhood-onset progressive myoclonic epilepsy [67] or familial dementia [68]. Amongst genes downregulated by exposure to VPA in our study was *Ap2b1*, which encodes an essential component of vesicle function, including synaptic vesicles [69]. These and other genes identified in this model system are novel candidates that may well be worthy of closer analysis in relation to the mood-stabilising effects of VPA.

In summary, using RNA-Seq analysis in a serotonergic cell culture model, we showed that therapeutically relevant doses of VPA and lithium resulted in differential regulation of a number of genes. Functional annotation revealed enrichment for several GO processes and canonical pathways that may be relevant to mood regulation. These include ECM remodelling, cell adhesion and chemotaxis. VPA resulted in the differential regulation of many more genes than lithium. Interestingly, both the genes regulated by lithium were also regulated by VPA. This suggests that both these drugs, although of quite different chemical composition, potentially act on overlapping genes and pathways to effect therapeutic outcomes.

This study on the transcriptional profile of a serotonergic cell line in response to the major mood stabilizers sheds some light on potentially novel molecular mechanisms underlying the action of these two drugs. This work also reveals potential regulatory processes that may occur in cells of the raphe nuclei after exposure to these drugs. Being buried deep in the brain, the serotonergic cell bodies of the raphe nuclei are not easily tractable for study. Further analysis of the genes and pathways identified in this work may provide novel insights into therapeutic mechanisms of actions for mood stabilizer drugs.

## Acknowledgement

This work was supported by the Jim and Mary Carney Charitable Trust, Whangarei, New Zealand. DB was supported by a Postgraduate Scholarship from University of Otago.

